# Dynamic roles of ILC3 in endometrial repair and regeneration

**DOI:** 10.1101/2024.08.10.606309

**Authors:** Antonia O. Cuff, Ee Von Woon, Thomas Bainton, Brendan Browne, Phoebe M. Kirkwood, Frances Collins, Douglas A. Gibson, Philippa T.K. Saunders, Andrew W. Horne, Mark R. Johnson, David A. MacIntyre, Victoria Male

## Abstract

Innate lymphoid cells (ILC) are prominent in the human uterine mucosa and play physiological roles in pregnancy. ILC3 are the second-most common ILC subset in the uterine mucosa, but their role remains unclear. Here we define two subsets of lineage-negative CD56+ CD117+ CRTH2-uterine ILC3, distinguished by their expression of CD127. The CD127-subset is most numerous and active during menstruation and immediately after parturition, suggesting a role in repair of the uterine mucosa (called endometrium outside of pregnancy); the CD127+ subset is most numerous and active immediately after menstruation, as the endometrium regenerates. In healthy endometrium, ILC3 are spatially associated with glandular epithelial and endothelial cells, which both express receptors for the ILC3-derived cytokines, IL-22 and IL-8. In the eutopic endometrium of people with endometriosis, ILC3 are located further from glandular epithelial and endothelial cells suggesting that these cells may be less exposed to ILC3 products, potentially with negative consequences for endometrial regeneration. Our findings highlight the dynamic nature of ILC3 in the uterine mucosa and suggest their primary role is in repair and regeneration. An improved understanding of uterine ILC3 will inform future research on endometrial health and disease.

## Introduction

Innate lymphoid cells (ILC) have essential roles at mucosal sites, contributing to barrier immunity and mucosal homeostasis. The ILC lineage broadly consists of three cell types: ILC1 (which is sometimes considered to include NK cells), ILC2 and ILC3.^1^ Uterine NK cells (uNK) are prominent in the human uterine mucosa, particularly in the first trimester of pregnancy, and are thought to have a role in placental implantation.^2^ ILC2 are not detectable in human uterine mucosa^2,3^ while ILC3 have been reported in the nonpregnant uterine mucosa (endometrium), and first and third trimester pregnant uterine mucosa (decidua).^3,4,5,6,7,8^ However, the functions of ILC3 in human reproduction are yet to be elucidated.

A major challenge in studying uterine ILC3 has been the lack of a consensus on their phenotype. In non-uterine tissues, ILC3 are most often identified by negative expression of markers of other immune cell lineages, and positive expression of CD127 (IL-7Rα, a pan-ILC marker) and CD117 (a marker of early lymphocyte development).^9,10^ In contrast to ILC3 in most other human tissues,^10^ uterine ILC3 express CD56.^ref.3,5,6,7^ Because of this, some studies of CD56+ cells that purportedly focus on uNK cells also include CD56+ uterine ILC3.^ref.11^

Human uterine ILC3 produce their prototypical cytokine IL22, as well as GM-CSF and IL-8.^ref.5,6,12^ IL-8 has been proposed to play a role in ILC3-neutrophil crosstalk which may be important for mediating transient inflammation associated with placental implantation.^12^ Moreover, it acts as a chemoattractant of placental extravillous trophoblast (EVT) cells, so could play a direct role in implantation.^13^ ILC3-derived IL-22 supports epithelial cell homeostasis, particularly in the presence of bacterial infection.^14^ Since trophoblast cells are epithelial, this suggests that ILC3 might also have a role in protecting against intrauterine infection during pregnancy. In support of this, IL-22 protects against LPS-induced pregnancy failure in mice,^15^ although during infection with the intracellular bacteria *Ureaplasma*, IL-22 was found to increase the rate of preterm birth.^16^

Here, we describe a method to identify ILC3 subsets in the human uterine mucosa by flow cytometry and high dimensionality reduction analysis, and use this to define the number and function of ILC3 subsets across the menstrual cycle, pregnancy and the postpartum period. We report that ILC3 number and activity is highly dynamic, peaking at times of endometrial repair and regeneration. We also find differences in ILC3 localisation in the eutopic endometrium of people with endometriosis, compared to those without endometriosis. Collectively, our findings suggest a primary role for ILC3 in endometrial repair and regeneration and potentially implicates these cells in the pathogenesis of endometriosis.

## Results

### Identification of ILC3 subsets in human uterine mucosa

Because there is currently no consensus definition of ILC3 across uterine mucosal tissues, cells of the uterine mucosa at different stages of the menstrual cycle and pregnancy (menses endometrium, 7; proliferative endometrium, 7; secretory endometrium, 7; first trimester decidua, 12; third trimester decidua, 8; postpartum endometrium, 5) together with matched peripheral blood cells were isolated and then analysed by flow cytometry (Figure 1a). We next examined CD45+ lineage negative (CD3, CD4, CD14, CD19, CD34) cells (Supplementary Figure 1) using an unsupervised approach. UMAP analysis revealed two major regions of cells: those originating from the uterine mucosa (grey) and those derived from the blood (red) (Figure 1b), consistent with previous reports.^7^ There was some overlap between uterine and peripheral blood ILC subsets, suggesting that some subsets share similar phenotypes in both compartments. An overrepresentation of cells expressing high levels of CD56 in the uterine mucosa was detected (Figure 1c), likely to include uterine NK cells, other CD56+ ILC1^ref.7^ and CD56+ ILC3.^ref.3,5,6^

**Fig. 1.**
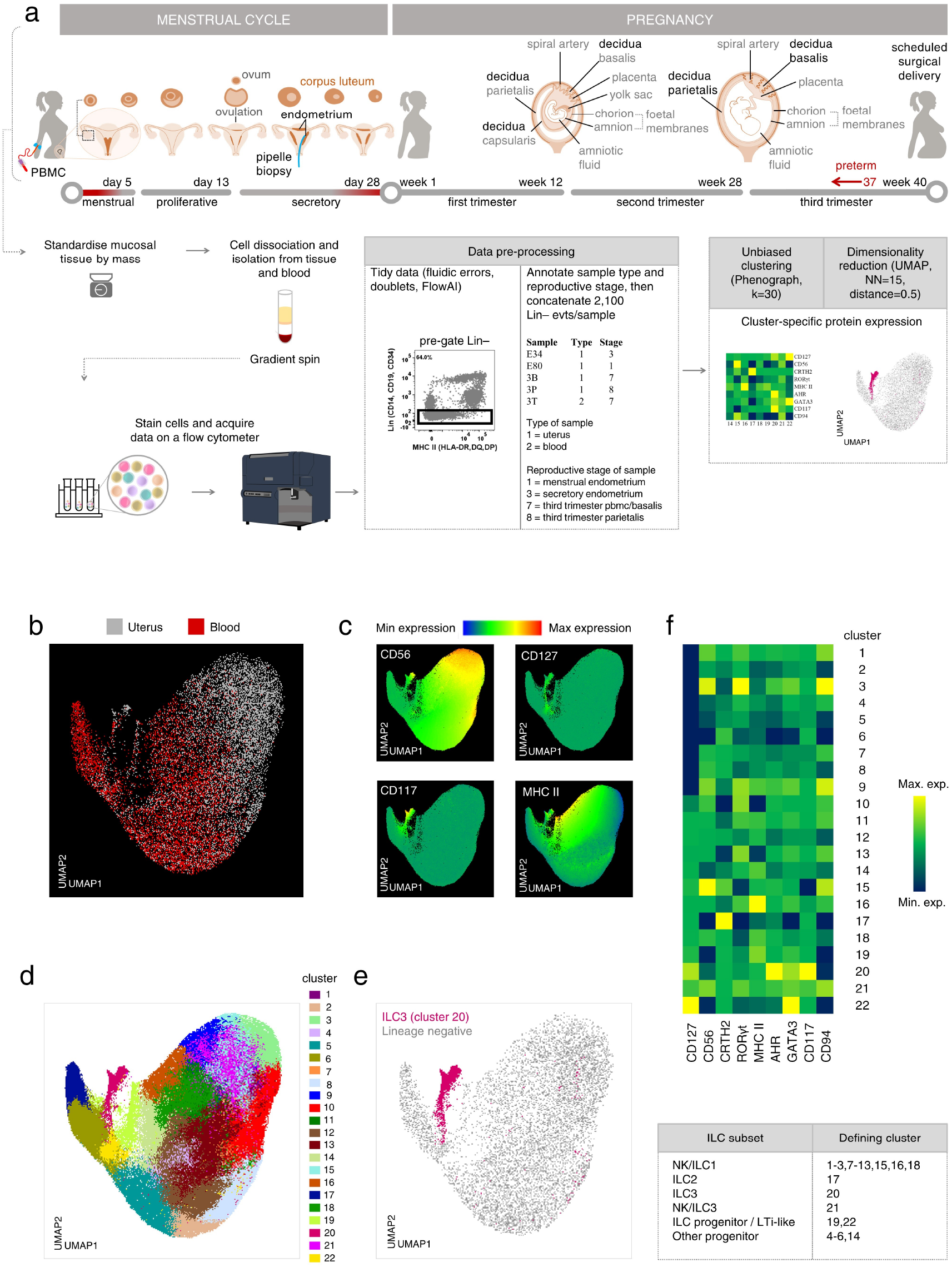
Identification of ILC3 subsets in human uterine mucosa. **A:** Schematic illustration of the stages of human reproduction, anatomical locations of specific types of uterine mucosa analysed (**bold** font) and workflow on uterine mucosal and matched peripheral blood samples. **B:** Single, live, CD45+ Lineage– (CD3, CD4, CD14, CD19, CD34) cells are distributed across the UMAP and coloured by origin (uterine mucosa, grey, 132,000 cells; blood, red, n = 108,000 cells). **C:** Relative intensity of expression of the indicated markers is shown. **D:** Unbiased clustering of CD45+ Lin– cells was completed using Phenograph. **E:** Cluster 20, with a phenotype consistent of ILC3, is highlighted in deep pink. **F:** The heatmap shows the geometric mean fluorescence intensities of the indicated markers in each cluster. Menses (M, n = 7), proliferative (P, n = 7), secretory (S, n = 7), first trimester (1T, n = 12), third trimester (3T blood, n = 8), decidua basalis (3B, n = 8) or parietalis (3P, n = 8), and postpartum (PP, n = 5).

To define ILC3, cells were clustered using Phenograph and those expressing ILC3 markers, including AHR, RORγt, CD117, CD127 and MHC II, ^ref.9,17^ were identified (Figure 1c). Cluster 20 expressed the ILC lineage-defining transcription factors AHR and RORγt, as well as high levels of CD117 and intermediate or bimodal levels of CD127. MHC II, associated with antigen presentation and immune cell activation, was expressed in cluster 20, as well as adjoining regions (Figure 1c-f). Identities were assigned to other clusters based on the expression of canonical markers, shown in Figure 1d,f.

We next used these features to design a conventional flow cytometric gating strategy to identify ILC3 in reproductive tissues. Lineage-negative CD94-CD127+ CD117+ CD56+, as previously described in decidua^5,6^ and secondary lymphoid tissues,^18^ were consistently identified in uterine mucosa spanning the menstrual cycle and pregnancy (Figure 2a). The lineage-defining transcription factors RORγt and AHR were expressed by these cells (Figure 2b). The bimodal distribution of CD127 in Cluster 20 also prompted us to examine lineage-negative CD94-CD127-CD117+ and CD56+ cells (Figure 2a). The expression of RORγt and AHR by these cells (Figure 2b) suggests that these are also ILC3: therefore, these CD127-ILC3 were also included in the downstream analyses of ILC3 dynamics in the human uterine mucosa.

**Fig. 2.**
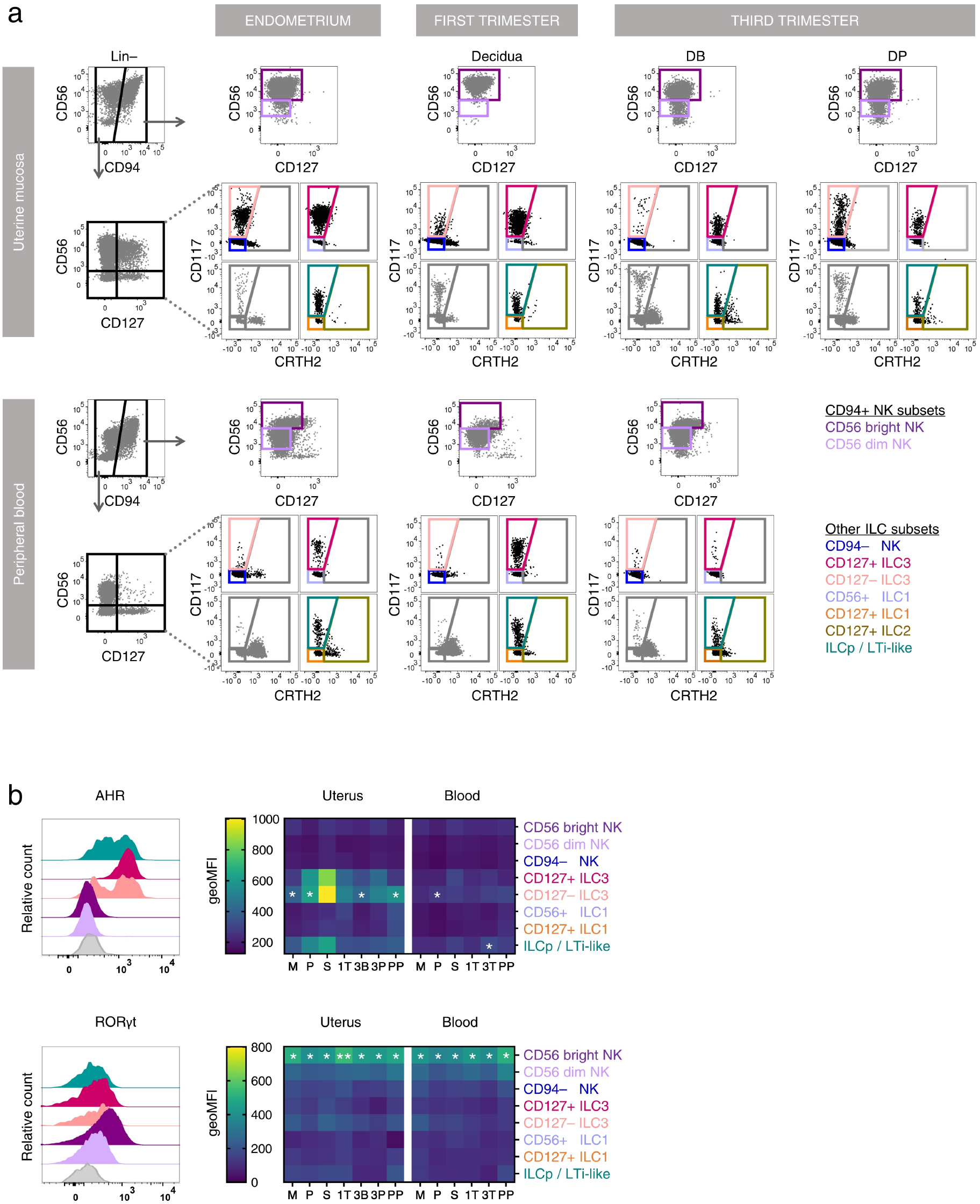
Uterine ILC3 are present throughout the menstrual cycle, pregnancy and post-partum. **A:** Flow cytometric gating strategy to identify ILC subsets among CD45+ Lin– immune cells isolated from uterine mucosal tissue and matched peripheral blood at different stages of the menstrual cycle, pregnancy and post-partum. **B:** Relative fluorescence intensities of staining for the ILC3 lineage-defining transcription factors, AHR and RORγt across the menstrual cycle and pregnancy. Menses (M, n = 7), proliferative (P, n = 7), secretory (S, n = 7), first trimester (1T, n = 12), third trimester (3T blood, n = 8), decidua basalis (3B, n = 8) or parietalis (3P, n = 8), and postpartum (PP, n = 5). Statistical significance was determined by normalization using Z score transformation (one-tailed, * p < 0.05, ** p < 0.01).

Lineage-negative CD94-CD127+ CD117+ CD56-cells are ILC progenitors (ILCp) in the blood^19^ but have also been called lymphoid tissue inducer (LTi)-like cells in the decidua.^5^ We identified cells with this phenotype in blood, decidua and endometrium (Figure 2a). As we did not assess either ILC progenitor or LTi potential of these cells, they are hereafter referred to as “ILCp/LTi-like cells”. These cells expressed AHR, but at lower levels than the ILC3 subsets (Figure 2b).

### Uterine ILC3 are most numerous and active at times of endometrial repair and regeneration

We next traced the distribution of ILC3 in uterine mucosal samples from healthy participants across reproductive stages. The frequency of uNK cells was used to validate immune cell isolation methods. Consistent with previous reports, their number and frequency peaked during the first trimester of pregnancy^20^ (Figure 3a,d).

**Fig. 3.**
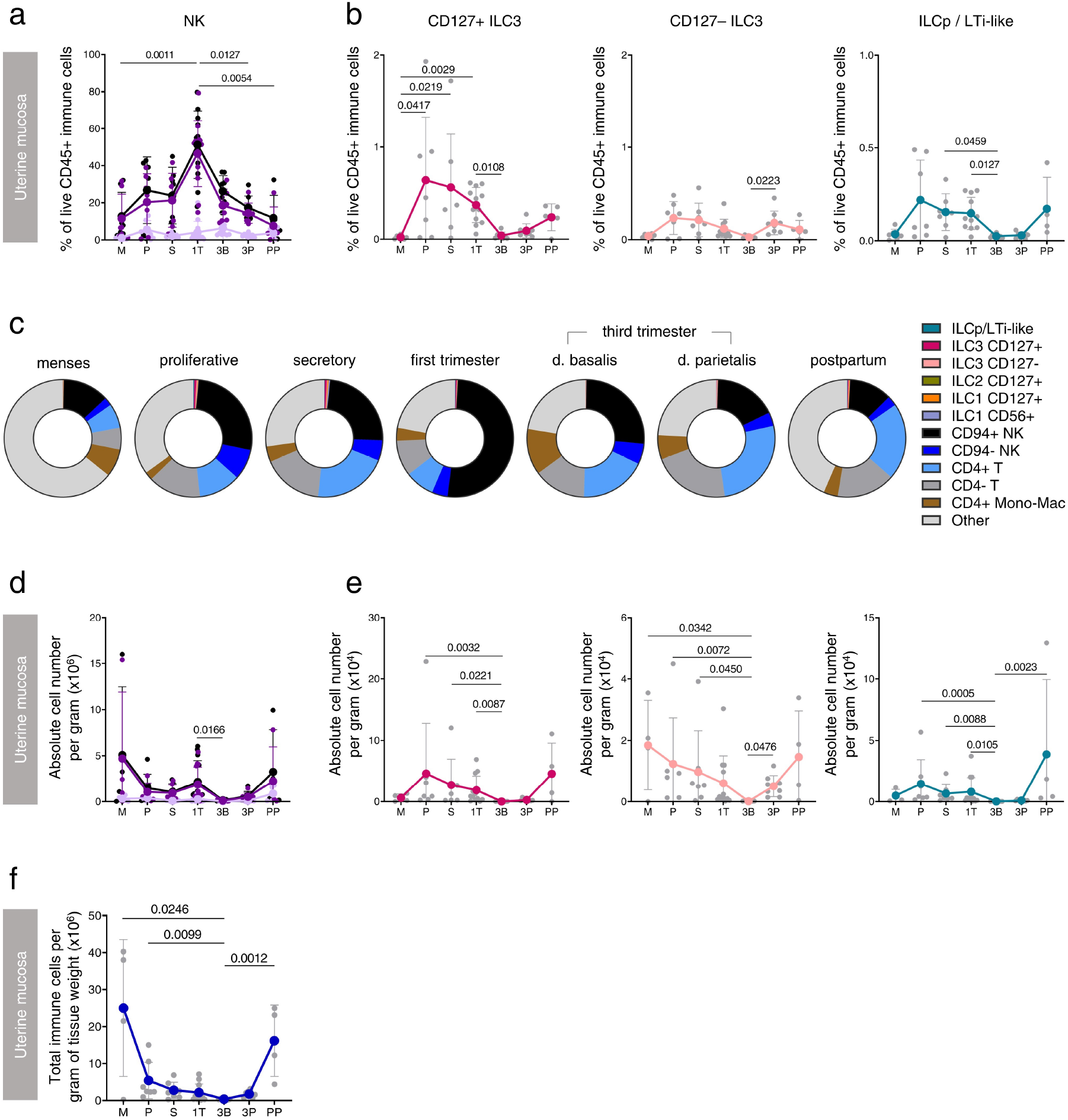
Uterine ILC are most numerous at times of tissue repair and regeneration. **A:** Frequency of NK cells in the uterus across the menstrual cycle and pregnancy. **B:** Frequency of ILC3 subsets in uterine mucosa across the menstrual cycle and pregnancy. **C:** Representative frequencies of all live CD45+ immune cell populations in the uterus at different stages of the human reproductive life cycle. **D:** Absolute number of NK cells per gram of tissue. **E:** Absolute number of ILC3 subsets per gram of tissue. **F:** Absolute number of all live CD45+ immune cells per gram of tissue. A Kruskal-Wallis statistical test for significance was performed with a Dunn’s correction for multiple hypotheses testing (* p < 0.05, ** p < 0.01). Menses (M, n = 7), proliferative (P, n = 7), secretory (S, n = 7), first trimester (1T, n = 12), third trimester (3T blood, n = 8), decidua basalis (3B, n = 8) or parietalis (3P, n = 8), and postpartum (PP, n = 5).

In contrast, CD127+ ILC3, CD127-ILC3 and ILCp/LTi-like cells were most frequent in the proliferative and secretory phases of menstrual cycle and in the post-partum period, together accounting for approximately 1% of all immune cells during their peak at proliferative phase of a menstrual cycle (Figure 3b,c). Cell frequencies can be due either to a decrease in their number or an increase in the number of other CD45+ cells (Figure 3c, Supplementary Figure 2), so we determined the absolute number of each subset per gram of tissue (Figure 3d,e). CD127+ ILC3 and ILCp/LTi-like cells were most numerous in regenerating proliferative and post-partum endometrium while CD127-ILC3 were most numerous during menstruation and in the post-partum period, times of endometrial repair. For all subsets, the fewest cells were observed during the third trimester of pregnancy. Fluctuations in the total number of immune cells are shown for context (Figure 3f). In the peripheral blood, few significant changes were observed, although there was a trend towards fewer CD127+ ILC3 and ILCp/LTi-like cells in the third trimester and post-partum (Supplementary Figure 3b,e).

We next examined the capacity of uterine ILC3 to produce IL-22 and IL-8. ILC3 subsets in the uterine mucosa produced more IL-22 and IL-8 than other ILC subsets, with CD127+ and CD127-ILC3, rather than ILCp/LTi-like cells, the major producers of these cytokines (Figure 4a,b). IL-22 was produced by uterine ILC3 subsets across all reproductive stages, but unstimulated CD127+ and CD127-ILC3 displayed significantly higher production of IL-22 during early menstrual cycle and post-partum phases. MHC II, which presents antigen and is a marker of immunological activation, was more highly expressed towards the end of pregnancy (Supplementary Figure 4). IL-8 production by CD127-ILC3 was also highest in the early menstrual cycle and immediately postpartum, although there was also some IL-8 production in the third trimester decidua basalis.

**Fig. 4.**
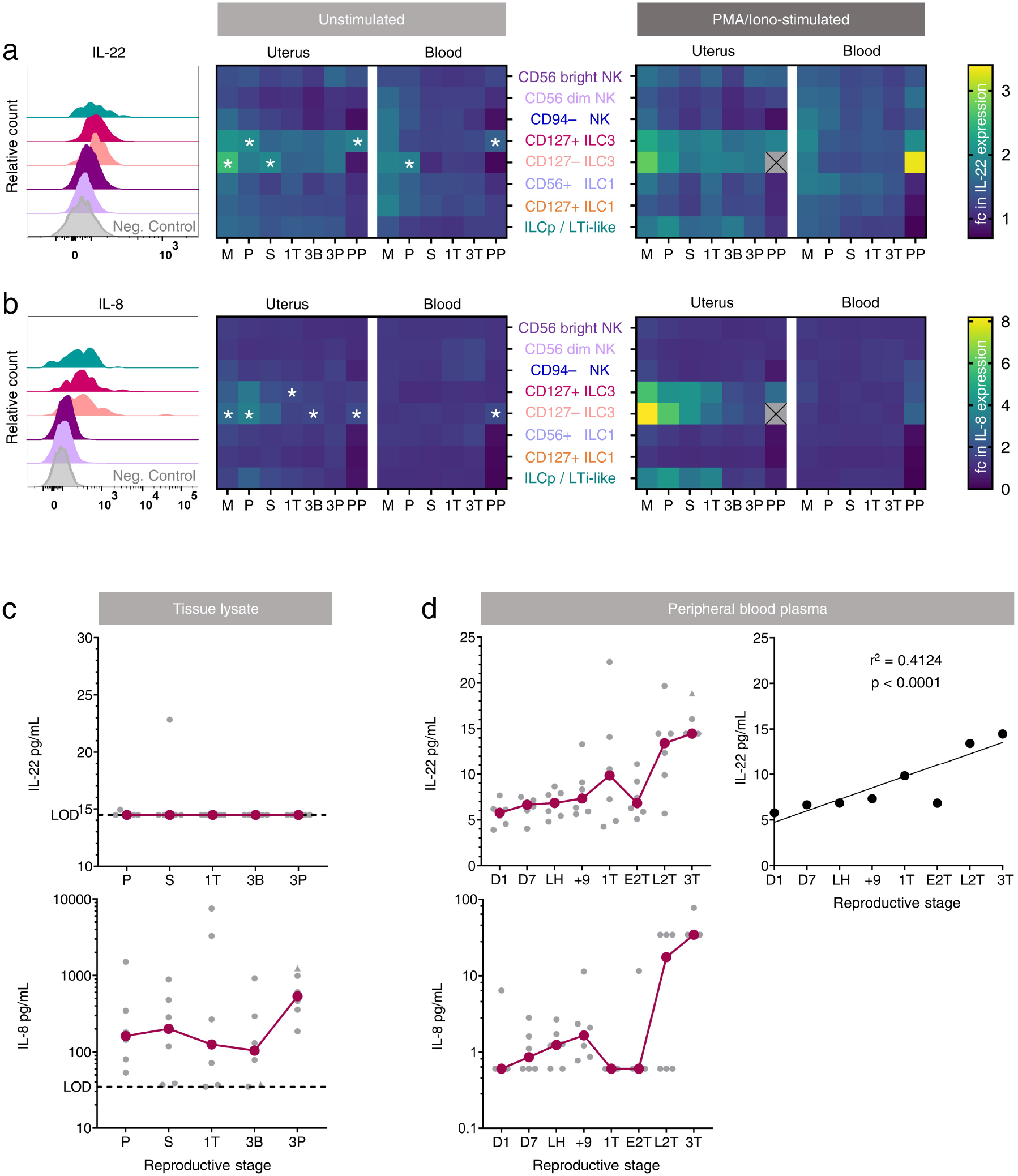
Uterine ILC produce IL-22 and IL-8 preferentially at times of tissue repair and regeneration. **A**,**B:** Isolated leukocytes were cultured directly ex vivo in the presence of Brefeldin A (10 μg/mL) and Monensin (2 μM) for 4 hours. Phorbol myristate acetate (PMA, 50 ng/mL) and Ionomycin (1 μg/mL) allowed the examination of effector molecule production by ILCs for the stimulation condition. Values are normalised to an internal negative control. Statistical significance was determined by Z score transformation (one-tailed, * p < 0.05). Histograms and heatmaps for IL-22 (**A**) and IL-8 (**B**) expression by ILC subsets across the menstrual cycle and pregnancy are shown. **C**,**D:** Profiling of IL-22 and IL-8 by bead-based array on uterine tissue lysates (**C**) and peripheral blood plasma (**D**). Linear regression between the level of plasma IL-22 and reproductive stage is shown. Stages examined included the menses (M, n = 5), proliferative (P, n = 6 uterus, n = 5 blood), secretory (S, n = 7), first trimester (1T, n = 11 uterus, n = 10 blood), third trimester (3T blood, n = 5), decidua basalis (3B, n = 5) or parietalis (3P, n = 5), and postpartum (PP, n = 3) stages. Peripheral blood plasma taken from menstrual cycle day 1 (D1), day 7 (D7), day of ovulation, identified by LH surge, 9 days post-ovulation, as well as peripheral blood plasma collected during the early and late second trimester of pregnancy were also examined. LOD, limit of detection.

To determine whether this cyclical production of IL-22 and IL-8 affected cytokine levels in the uterine mucosa, we examined total IL-22 and IL-8 levels in tissue lysate (Figure 4c). IL-22 was generally below the threshold of detection, but intra-tissue levels of IL-8 were discernible and remained constant over reproductive stages. By contrast, IL-22 was detectable in plasma, with an increasing trend across progressive reproductive stages beginning from the first day of the menstrual cycle and ending at the non-labouring third trimester (Figure 4d). IL-8 was also largely detectable in plasma, although notably not during the first and early second trimester in this cohort.

### IL-22 and IL-8 receptors are expressed by epithelial and endothelial cells

To determine which cell types could be responding to ILC3-derived IL-22 and IL-8, we next examined the expression of the receptors for these cytokines. The expression of the IL-22 receptor (a dimer of IL10R2 and IL22R1) and IL-8 receptors CXCR1 and CXCR2 was examined on CD45-cell subsets identified as shown in Supplementary Figure 5. The expression of each receptor was normalised to the fluorescence minus one (FMO) control (Figure 5a).

**Fig. 5.**
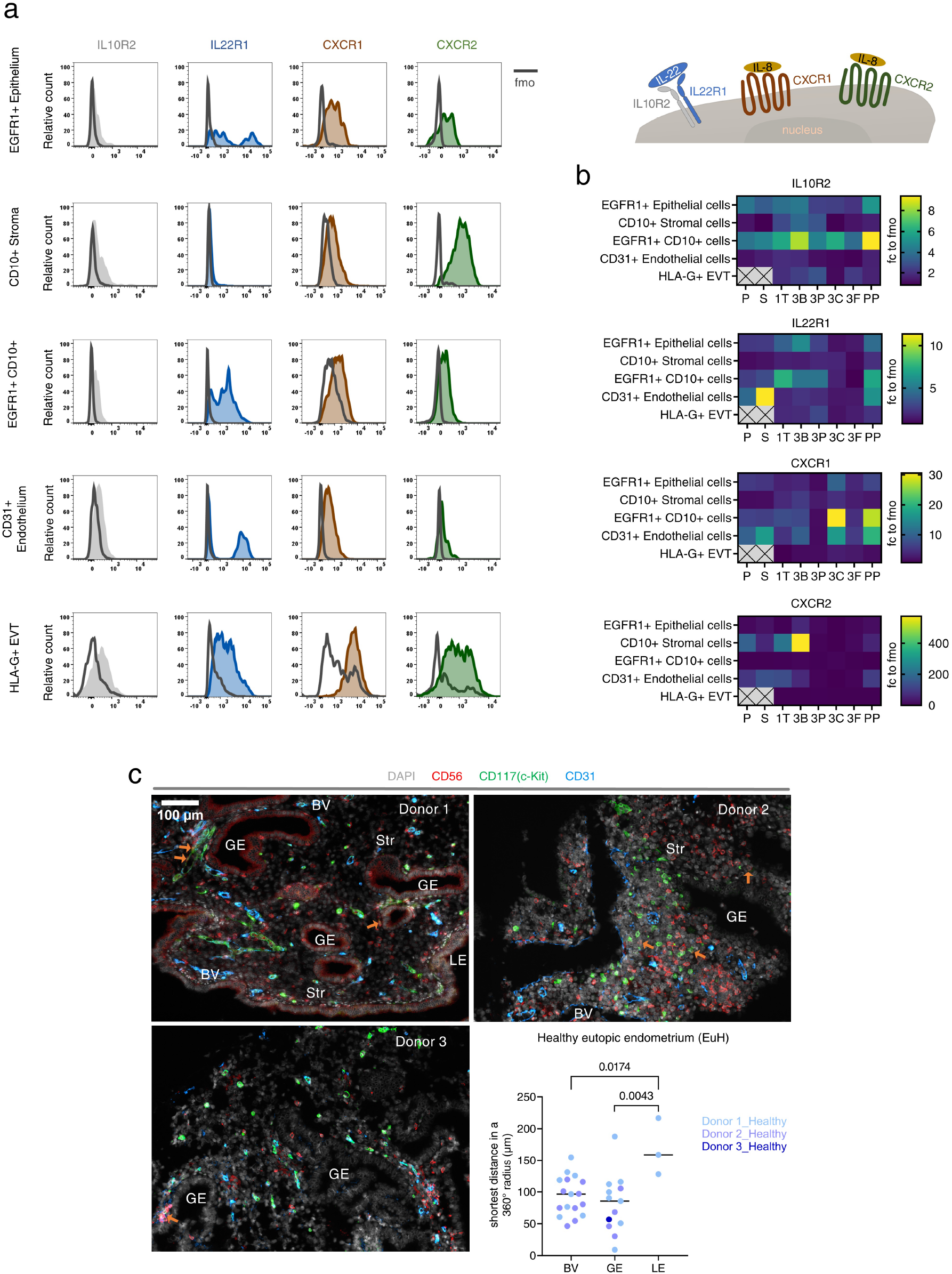
ILC3 are located near endothelial and glandular epithelial cells, which express IL-22 and IL-8 receptors. **A:** Representative histograms of IL10R2, IL22R1, CXCR1, and CXCR2 relative to the fluorescence minus one (FMO) control in different cellular components of the uterine mucosa. **B:** Summarised heatmap of receptor expression, normalised to the FMO control. Proliferative (P, n = 2), secretory (S, n = 4), (1T, n = 10), third trimester decidua basalis (3B, n = 7), decidua parietalis (3P, n = 7), chorionic membrane (3C, n = 2), foetal placenta (3F, n = 2) and postpartum (PP, n = 2). **C:** Spatial location of ILC3 in the functional layer of endometrium from three healthy donors are shown through confocal microscopy. Markers included CD56 (red), CD117 (green), CD31 (blue) and DAPI staining for the nucleus (grey). Orange arrows pinpoint ILC3, defined as CD56+ CD117+. The distance between each ILC3 and their surrounding endometrial structural component were determined using Fiji and statistically significant differences are indicated (one-way nested ANOVA, two-tailed).

Of the two IL-22 receptor subunits, IL10R2 was more widely expressed than IL22R1 across reproductive stages (Figure 5b). EGFR1+CD10-epithelial cells and EGFR1+CD10+ cells highly express both IL10R2 and IL22R1. CD31+ vascular endothelial cells uniformly express a low level of IL10R2 while a subset also express IL22R1. During pregnancy, EVT express both subunits of the IL-22 receptor, albeit at a low level compared to the FMO control.

Some expression of the IL-8 receptors CXCR1 and CXCR2 was observed across all cell types examined (Figure 5a) but their expression was more variable across reproductive stages than that of the IL-22 receptor subunits (Figure 5b). The highest levels of CXCR1 expression were observed in CD31+ vascular endothelial and EGFR1+CD10+ cells isolated from third trimester chorionic membrane, and from post-partum endometrium. In contrast, CXCR2 was more highly expressed in CD10+ stromal cells, particularly those isolated from third trimester decidua basalis.

### Uterine ILC3 are located near blood vessels and glandular epithelium

To investigate the propensity for interaction between uterine ILC3 and these non-immune cell types, we next sought to establish their proximity to each other. We focused on the endometrium since ILC3 were most numerous and active at this reproductive stage, examining endometrial samples from donors who, on investigation, were not found to have a gynaecological pathology (hereafter referred to as “healthy donors”). Multiplex immunofluorescence microscopy showed that ILC3 markers such as RORγt, AHR and CD127, which have been previously used on frozen tissue,^21,22,23^ were masked by fixation. Therefore, we identified ILC3 as CD56+CD117+, indicated by orange arrows (Figure 5c). The majority of CD56+CD117-cells represent uNK cells, although this group may also include some CD56+ innate T cells. Vascular endothelial cells (labelled BV) were identified by staining for CD31. Other cells were identified by morphology: luminal epithelial cells (a wall of columnar epithelia; LE), glandular epithelial cells (a mixture of cells with epithelial cell morphology largely gathered around a central cavity; GE) and stromal cells (the cell network which makes up the bulk of the endometrial structure and lies between the glands and blood vessels; Str).

CD56+ CD117+ ILC3 were found in the endometrium of all three healthy donors examined, although they were less frequent than CD56+ CD117-uNK cells, consistent with our observations by flow cytometry. ILC3 were located within the stroma, but close to blood vessels and glands. Quantification of the shortest distance between ILC3 and different endometrial components revealed that ILC3 were located at approximately equivalent distances from blood vessels and glands, but significantly further away from the luminal epithelium (Figure 5c).

### ILC3 are located further from blood vessels and glandular epithelium in endometriosis

Our observation that ILC3 are most abundant and active at times of endometrial regeneration suggested a role in this process. Further, our findings that ILC3 are most closely associated with blood vessels and glands, and that vascular endothelial and epithelial cells express receptors for their products, are consistent with interactions between ILC3 and these cells. Therefore, we sought to investigate whether the dynamics and localisation of ILC3 differed between healthy donors and those suffering from menstrual cycle disorders. We focused on endometriosis because of the well-established role of immune dysregulation in the pathology of the disease,^24,25^ (Supplementary Figure 6), including a recent report that ILCs are reduced in the endometrium during endometriosis.^26^

CD127+ ILC3 in the eutopic endometrium of donors with endometriosis (EuE) were less frequent than in eutopic endometrium from healthy donors (EuH), although the difference was not significant, and there was no difference in the frequency of CD127– ILC3 (Figure 6a). In donors with endometriosis, the frequency of both ILC3 subtypes in the ectopic endometrium was comparable to their frequency in eutopic endometrium. ILCp/LTi-like cells tended towards a higher frequency in eutopic endometrium of donors with endometriosis than in eutopic endometrium from healthy donors, although this difference was not significant. Peripheral blood ILC3 subtypes did not differ in donors with endometriosis relative to equivalent cells in healthy donors (Figure 6b).

**Fig. 6.**
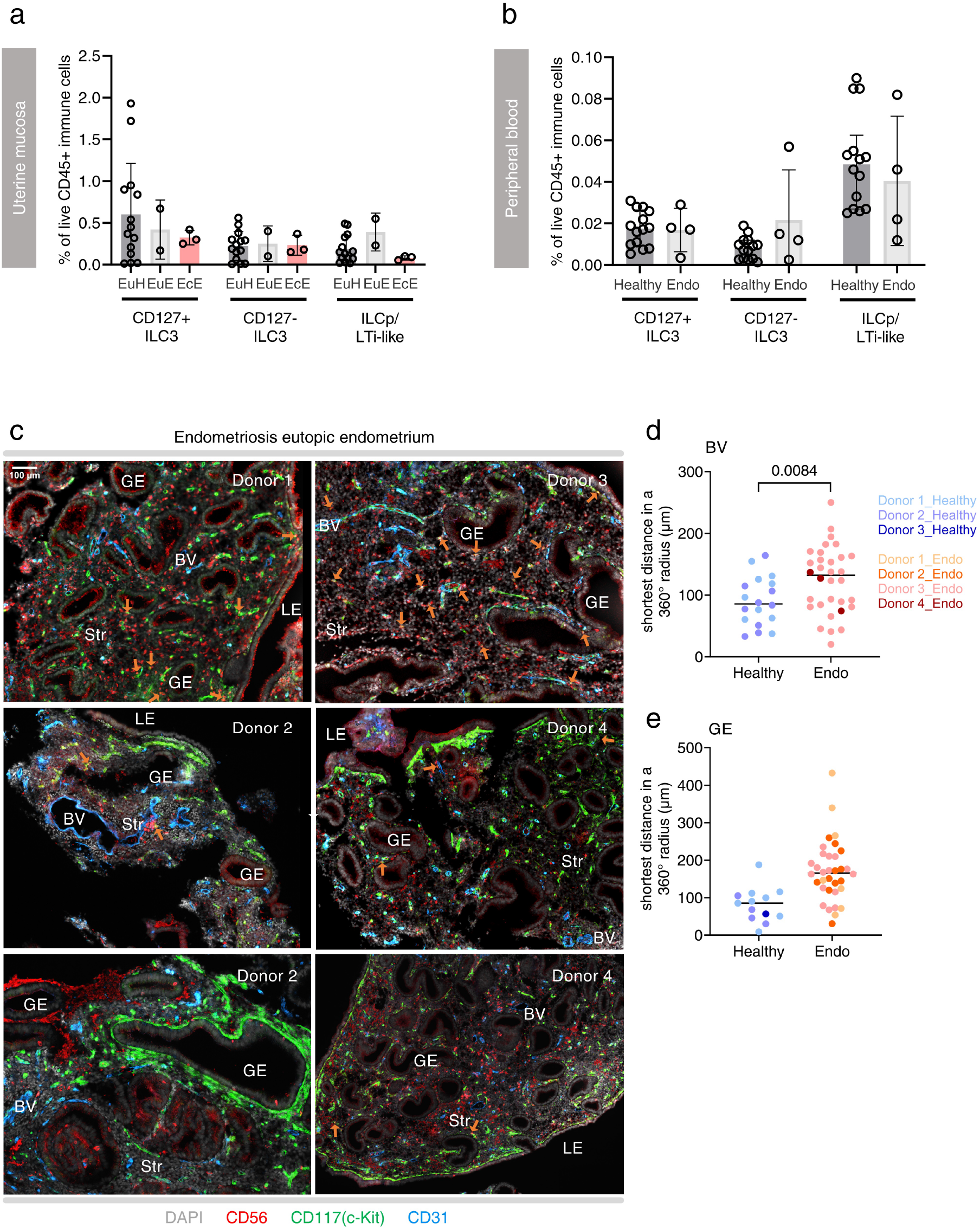
Endometrial ILC3 are located further from endothelial and glandular cells in endometriosis. **A:** Frequency of ILC3 subsets among total live CD45+ immune leukocytes isolated from eutopic endometrium in healthy donors (EuH, proliferative (P, n = 7), secretory (S, n = 7), eutopic endometrium in donors with endometriosis (EuE, proliferative (P, n = 1), secretory (S, n = 1) and ectopic endometrium in donors with endometriosis (EcE, proliferative (P, n = 1), secretory (S, n = 1), undefined (n = 1). **B:** Frequency of ILC3 subsets in the blood of healthy donors and donors with endometriosis (Healthy donors, proliferative (P, n = 7), secretory (S, n = 7; Endometriosis donors, proliferative (P, n = 1), secretory (S, n = 2), undefined (n = 1). **C:** Spatial location of ILC3 in the functional layer of eutopic endometrium from four donors with endometriosis are shown through confocal microscopy. Markers included CD56 (red), CD117 (green), CD31 (blue) and DAPI staining for the nucleus (grey). The distance between each ILC3 and their surrounding endometrial structural component were determined using Fiji and statistically significant differences are indicated (nested t test, two-tailed).

**Fig. 7.**
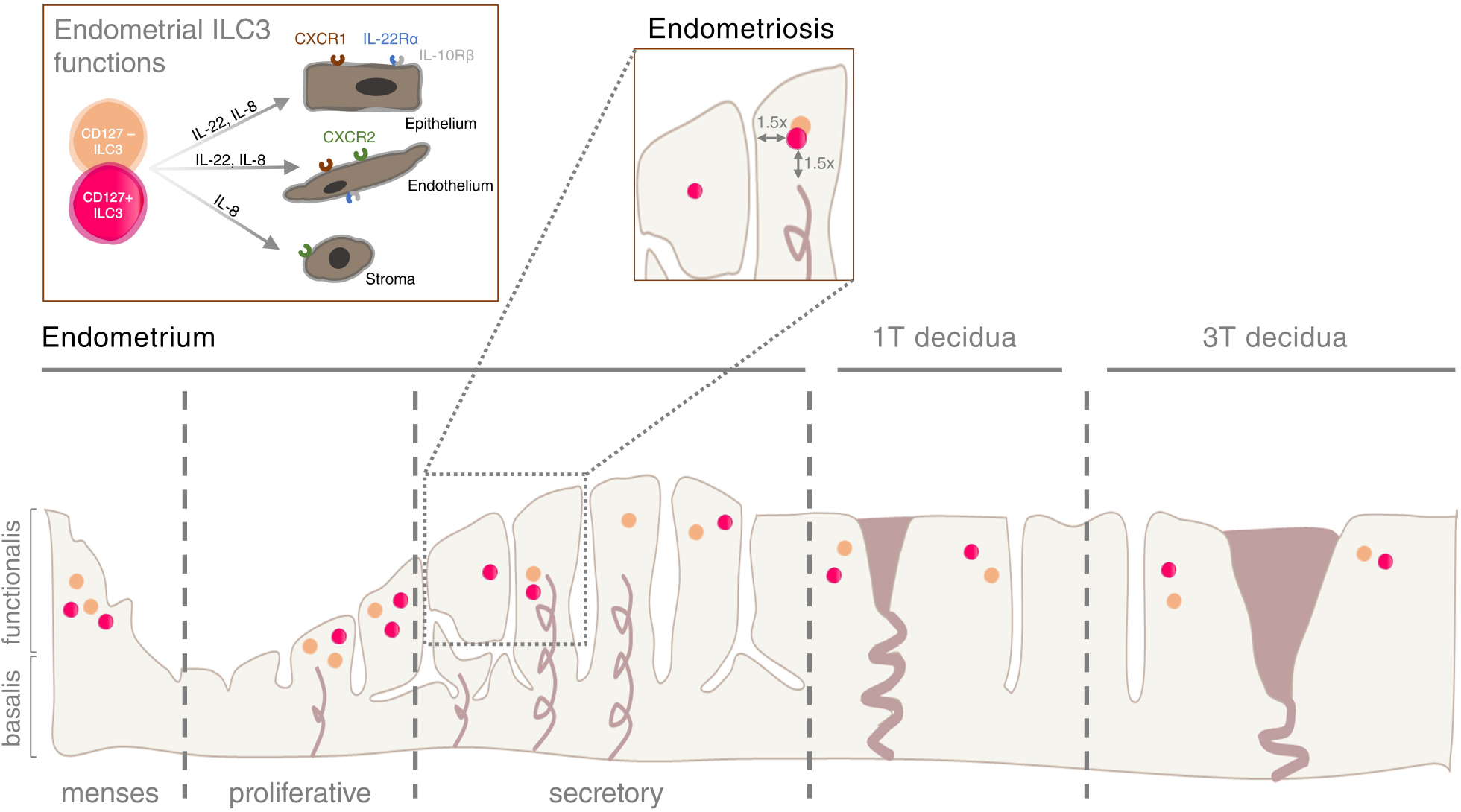
ILC3 during the healthy reproductive cycle and endometriosis. The prominence of uterine ILC3 at times of endometrial repair and regeneration implies that this may be their primary function. During the healthy menstrual cycle, endometrial ILC3 may regulate endometrial processes by secreting effector molecules, including IL-8 and IL-22, that target receptors which are differentially expressed by the epithelium, endothelium and stroma. While ILC3 frequencies are not altered during endometriosis, ILC3 are located further away from the endothelium and glandular epithelium and endometrial processes during endometriosis may change because of ILC3 displacement.

Next, we used multiplex immunofluorescence microscopy to examine the spatial distribution of ILC3 in the eutopic endometrium of donors with endometriosis compared to that of healthy donors (Figure 5c). In line with our findings by flow cytometry, ILC3 were also detected in the eutopic endometrium of endometriosis patients by microscopy (Figure 6c). These were located further away from blood vessels and glands than in controls, although the difference was only significant for blood vessels (Figure 6d,e).

## Discussion

Group 1 ILCs in the form of uNK cells are well represented in the human uterine mucosa, while ILC2 are rare. Here we provide new insight into the dynamics of ILC3 across the menstrual cycle and pregnancy. Previous attempts to define the dynamics and behaviour of ILC3 in the human uterus have been hampered by the lack of a consistent method to identify these cells, as well as the extent to which ILC3 share markers with more prominent uNK cell populations. Using high dimensionality reduction analysis, we identified two ILC3 subsets in the human uterine mucosa, both of which are detectable throughout the menstrual cycle and pregnancy. The ILC3 identity of the subsets was confirmed by their expression of the lineage-defining transcription factors AHR and RORγt and their production of the prototypical ILC3 cytokine IL-22. In line with a previous report that CD56^bright^ NK cells express RORγt in secondary lymphoid tissue,^27^ we also observed RORγt expression in CD56^bright^ uNK cells. Of note, one of the ILC3 subsets we identified expressed little of the canonical ILC surface marker, CD127. CD127-ILC2 have recently been reported^28^ but to our knowledge, this is the first such description of CD127-ILC3.

Previous analyses of group 1 uterine ILC frequencies found that peak uNK cell frequency and function coincided with the physiological processes with which they are most closely associated, implantation and placentation.^20,29^ Equipped with the ability to discern ILC3 subsets in the human uterine mucosa, we took a similar approach to determine the stage at which ILC3 are most active, and therefore when they are most likely to have a physiological role. One challenge of this approach is that the uterine mucosa is highly dynamic, with cyclic changes in other immune cell subsets impacting any analysis based only on cell frequencies: to mitigate this we also examined absolute changes in cell number per gram of tissue. Based on previous work in mice^15,30^ and humans^12^ we expected that ILC3 might have a role in pregnancy. However, CD127-ILC3 were most numerous and produced the most IL-22 and IL-8 at times of endometrial repair, during menstruation and in the immediate post-partum period. CD127+ ILC3 were most numerous and produced the most IL-22 in proliferative and post-partum endometrium, suggestive of a role in endometrial regeneration.

Further to analysing ILC3 production of IL-22 and IL-8, we also looked at the propensity for target cells in the uterine mucosa to respond to these cytokines. In line with a previous study by immunohistochemistry,^31^ we found IL10R2 expression in the epithelial and stromal cell compartments, as well as in EGFR1+CD10+ cells, which may represent stromal cells undergoing mesenchymal to epithelial transformation during endometrial repair.^32^ IL22R1 was also expressed by these cells throughout the menstrual cycle, in contrast to the findings of an earlier immunohistochemistry-based study that detected IL22R1 only in secretory endometrium.^31^ In our study, endothelial cells in the same stages prominently expressed IL22R1 but only expressed IL10R2 at low levels. We detected expression of the IL-8 receptors CXCR1 and CXCR2 among endothelial, stromal and stromal-epithelial intermediate cells, but little expression among epithelial cells. This is consistent with a previous study which detected CXCR1 and CXCR2 in endometrial stromal cells by immunohistochemistry, although this study also reported no expression of these receptors in endothelial cells, in contrast to our findings.^33^ The differences between our findings and those previously reported may be accounted for by the fact that flow cytometry enables the analysis of a larger number of cells and is thus better able to detect small numbers of highly expressing endothelial cells.

Our analysis of the spatial distribution of ILC3 in the endometrium finds that they are more closely associated with blood vessels and glands than luminal epithelium. This supports the concept that ILC3 are most abundant deeper in the uterine mucosa, similar to their location at other mucosal sites.^34^ Expression of receptors for ILC3 products by epithelial and endothelial cells, and the proximity between ILC3 and these cells is indicative of crosstalk between ILC3, endometrial glands and blood vessels.

To investigate this further, we characterised the dynamics and localisation of ILC3 in the endometrium of donors with endometriosis, compared to healthy control donors. It has recently been reported that the proportion of CD127+ ILC3 is reduced in the endometrium of donors with endometriosis, compared to healthy donors, while ILC3 proportion in blood is unchanged.^26^ Similarly, we also found CD127+ ILC3 less frequent in the eutopic endometrium of donors with endometriosis compared to healthy donors, although this difference was not statistically significant in our cohort.

ILC3 in the endometrium from donors with endometriosis were located significantly further from the vasculature than those in controls and a similar trend was observed for ILC3 location compared to the glands. This raises the possibility that endothelial and glandular epithelial cells are less exposed to ILC3 products in donors with endometriosis. IL-22 is well-known to promote epithelial cell proliferation^35^ and there is also some evidence it can act on endothelial cells to promote angiogenesis.^36^ Both processes are important for repair of the uterine mucosa, which occurs simultaneously with menstruation, and subsequent regeneration. Of note, people with endometriosis are approximately five times more likely to report heavy menstrual bleeding than controls.^37^ Although it has been suggested that the comorbidity between endometriosis and heavy menstrual bleeding results from progesterone resistance and endometrial hyper-plasia, this view has become increasingly contested^38^ and there is some evidence that the major endometrial ILC subset, uNK cells, are dysregulated in heavy menstrual bleeding.^39^ Our results are consistent with the concept that heavy menstrual bleeding in endometriosis could in part result from inadequate endometrial repair. Further studies determining whether other etiologies of heavy menstrual bleeding are also associated with differential distribution of endometrial ILC3 are warranted.

While our study reveals important new insight into ILC3 function in the uterine mucosa, it was limited to the examination of the upper layer of the endometrium (functionalis). Location of ILC3 in the deeper layer (basalis) remains to be determined. Further, while the prominence of ILC3 in the non-pregnant endometrium suggests a physiological role outside of pregnancy, we do not rule out a role in pregnancy. Expression of IL-22 and IL-8 receptors by extravillous trophoblasts indicates that they stand ready to detect ILC3 products if they are produced, perhaps in pathological situations. Our finding that ILC3 express MHC II more highly towards the end of pregnancy could also imply that ILC3 may have roles other than IL-22 production at this time. Interactions of ILC3 with other immune cells in the uterus akin to mucosal tissues elsewhere also remains to be elucidated. These include their ability to present antigens to T cells^17,21,40,41,42^ and regulate myeloid cell functions^43^ through effector molecules released into the local tissue microenvironment, mechanisms that will amplify the impact of this small immune cell population.

In summary, our data indicate a primary role for ILC3 in endometrial repair and regeneration and suggest that they may play a role in the pathogenesis of endometriosis. Furthermore, our comprehensive description of the dynamics and localisation of uterine ILC3 across the healthy menstrual cycle and pregnancy provides a reference from which their role in gynaecological and obstetric pathologies can now be explored.

## Methods

### Human tissue

The collection of fresh human tissue was approved by London Chelsea Research Ethics Committee (study numbers 10/H0801/45 for endometrium and 11/LO/0971 for decidua). Samples of the uterine mucosa were collected at different stages of the menstrual cycle and pregnancy (menses endometrium; proliferative endometrium; secretory endometrium; first trimester decidua; third trimester decidua; postpartum endometrium) together with matched peripheral blood samples. Functionalis endometrium was collected as pipelle biopsies from healthy donors with regular cycles at scheduled clinics. Menstrual cycle phase was confirmed by donor’s self-reported cycle day in combination with serum progesterone levels against a reference dataset.^44^ Decidual samples from weeks 6 – 15 were obtained during pregnancy terminations upon personal request by the donor. Non-labouring decidual samples in third trimester pregnancies were collected at term (weeks 37 – 40) during elective caesareans. Pipelle en-dometrial biopsies were collected from donors at the time of surgery, formalin-fixed and embedded (Lothian Research Ethics Committee 11/AL/0376). Donor demographics are shown in Supplementary Tables 1 – 4.

Fresh pipelle endometrial biopsies were mechanically disaggregated through a 100 μm cell strainer, centrifuged (700xg, 10 min, RT) and resuspended in 10% FBS-supplemented PBS. Filtrates underwent repeat filtration though a 70 μm strainer and leukocytes were separated by density gradient centrifugation over Histopaque (700xg, 20 min, RT without brake).

First trimester decidual samples were processed as described.^20^ Briefly, tissues were rinsed in PBS to remove excess blood and most of the placenta was removed by excision. Using scalpel blades, decidual tissues were minced then homogenised in 10% FBS-supplemented PBS on a GentleMACS processor (Miltenyi). The homogenate underwent further mechanical disaggregation through a 75 μm sieve with regular washes in PBS. Filtrates were centrifuged (500xg, 10 min, RT), resuspended in 10%-FBS supplemented PBS and underwent density gradient centrifugation over Histopaque (700xg, 20 min, RT, without brake).

Decidua basalis and parietalis were dissected from term placentae as described.^45^ To isolate leukocytes and placental cells, tissues were mechanically disaggregated by GentleMACS homogenisation in 1ml Accutase per gram tissue. Homogenised tissue was then digested (100 rpm, 1 h, 37°C), digests were filtered through a 70 μm strainer and leukocytes separated by density gradient centrifugation over Histopaque (700xg, 20 min, RT, without brake). Non-immune cells other than placental cells were isolated as described.^46^ Dissected tissues were cleansed of blood in PBS and homogenised in 0.25% trypsin-enriched PBS containing 0.05% DNAse I. Tissues were enzymatically disaggregated (100 rpm, 15 min, 37°C) before trypsin activity was quenched in serum-rich media (10% Newborn Calf Serum-supplemented DMEM/F12 medium, plus 1% Penicillin-Streptomycin and 0.4% DNase I). Digests were filtered through a gauze mesh, centrifuged (500 rpm, 20 min, RT), resuspended in serum-rich media, refiltered through a 100 μm strainer and overlayed on Histopaque for cell isolation by density gradient centrifugation (700 xg, 20 min, without brake).

Tissue lysates were prepared in PBS supplemented with protease inhibitors using a Precellys homogeniser system (8μL per mg tissue > 60 mg, or 5μL per mg tissue < 60 mg). Lysates were allowed to chill at 4°C before final centrifugation at 16,000xg, 10 min, 4°C for collection. A Bio-Rad DC protein assay was used to quantify protein content before storing lysates at -80°C.

### Flow cytometry

Viable cells were identified using fixable viability stain eFluor 450 (eBioscience) in PBS (15 min, 4°C). Non-specific antibody staining was inhibited using blocking buffer (male 10% human AB serum, 1% bovine serum albumin, 2 mM EDTA, 25 μg BD Fc Block; 15 min, 4°C) and washed before completing surface stain in staining buffer (PBS supplemented with 1% FBS, 2mM EDTA). Brilliant stain buffer (BD Biosciences) was added to the surface stains to counteract anticipated intra-brilliant violet fluorescence interferences. Intracellular antigens were stained using the Human Foxp3 Fix/Perm buffer (BD Biosciences). Surface-stained leukocytes were fixed (10 min, RT), washed then permeabilised (30 min, RT) before intracellular staining antibodies (Supplementary Table 5) were added (30 min, RT). Washes at the end of each step were completed in staining buffer. Data were acquired on a BD LSR Fortessa X-20 cytometer. Details of antibodies are given in Supplementary Table 5.

Immune cell frequencies and mean fluorescence intensities were extracted using FlowJo (Treestar). Normalised immune cell counts were generated by multiplying the frequency of each cell type (as a proportion of live CD45+ cells) by the absolute number of immune cells isolated per gram of tissue. High dimensionality reduction analyses were also completed in FlowJo, following data preparation in line with the current recommendations for optimal cytometry experiments, eliminating anomalies such as fluidic errors, doublets, and dead cells.^47,48^

### Functional assays

For staining of effector molecules, cells were cultured for 4 hours in RPMI-supplemented media (RPMI 1640 – Glutamax, 10% Foetal Bovine Serum, Penicillin-Streptomycin, 1x Non-essential amino acids, 1mM Sodium pyruvate, 25mM HEPES, 50μM B-mercaptoethanol), and in the presence of Brefeldin (10μg/mL) and Monensin (2μM). Cells were cultured either without stimulation or with a non-specific stimulation with Phorbol myristate acetate (PMA, 50ng/mL, Sigma) and Ionomycin (1μg/mL, Sigma).

### Immunofluorescence microscopy

Endometrial tissue sections (5 μm thick) were dewaxed in xylene and rehydrated in serial ethanol dilutions. Antigens were unmasked by sequential antigen retrieval as in Supplementary Table 6a. Tissues were permeabilised in 0.25% Triton in PBS (30 min, RT). Non-specific antigen detection was achieved by incubation in 3% hydrogen peroxide (30 min, RT) and serum-rich block (5% normal donkey serum, 2% bovine serum albumin in PBS, 30 minutes, RT). Primary antibodies (Supplementary Table 6a) attached overnight at 4°C and after washes in 0.05% Tween-20-supplemented PBS were detected using HRP-conjugated secondary antibodies (Supplementary Table 6b) and HRP-activated dyes (Supplementary Table 6c). Tissues were mounted in DAPI and imaged. Images were captured on a Zeiss Axioscan Z1 fluorescence microscope, processed in ZEN Blue and processed images were analysed in FIJI.

### Multiparametric bead-based immunoassays

IL-8 and IL-22 in plasma and tissue lysate were measured using a custom Legendplex (Biolegend) panel targeting TNF-β (A3), IL-1β (A4), IFN-γ (A6), TNF-α (A10), IL-8 (B2), IL-17A (B4), GM-CSF (B5), IL-23 (B7), IL-22 (B9).

### Statistical analysis

Data were tested for normality using Shapiro Wilk’s test, to inform downstream choice of statistical testing. GraphPad Prism was used to plot data and perform statical tests. Details of the specific test used in each case are given in the corresponding figure legends.

## ACKNOWLEDGEMENTS

This study was funded by Borne. We would like to thank all the people from Chelsea and Westminster Hospital, West Middlesex University Hospital, and John Hunter Clinic (London, UK), who contributed samples to this study.

## Data availability

The data underlying this publication has been deposited at https://osf.io/c439u/

