## Supplementary_Figures for "Dynamic roles of ILC3 in endometrial repair and regeneration"

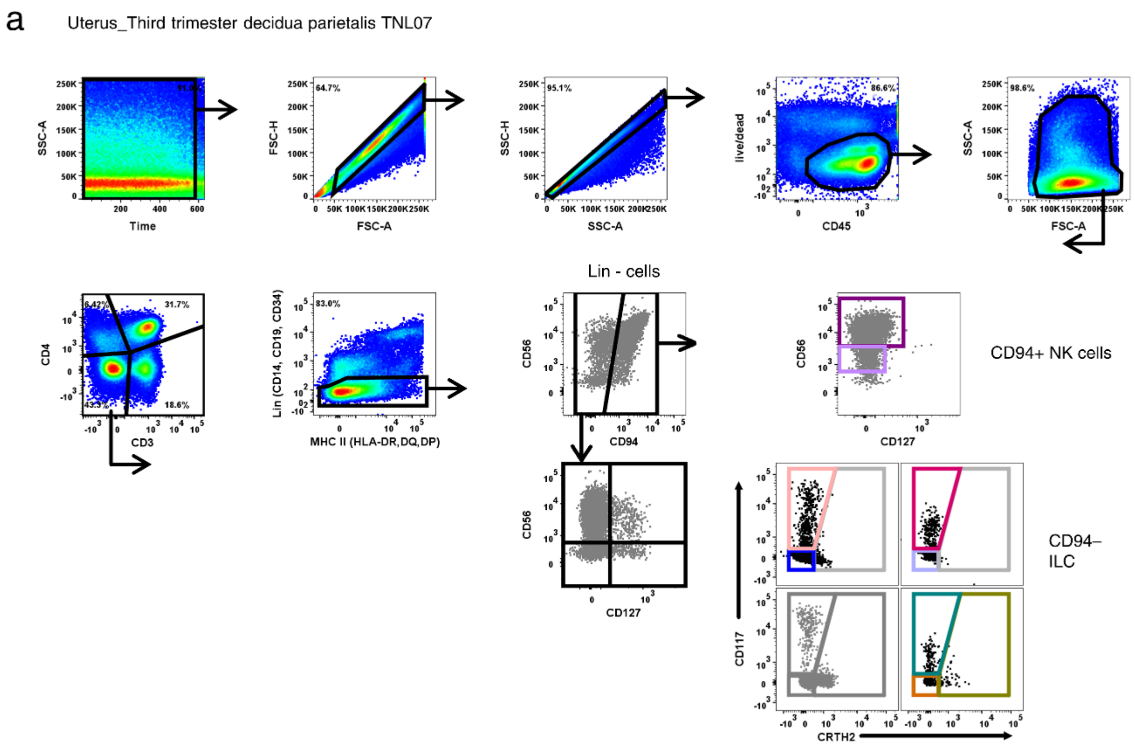


**Supplementary Figure 1. a** Complete flow cytometric gating strategy for ILCs.

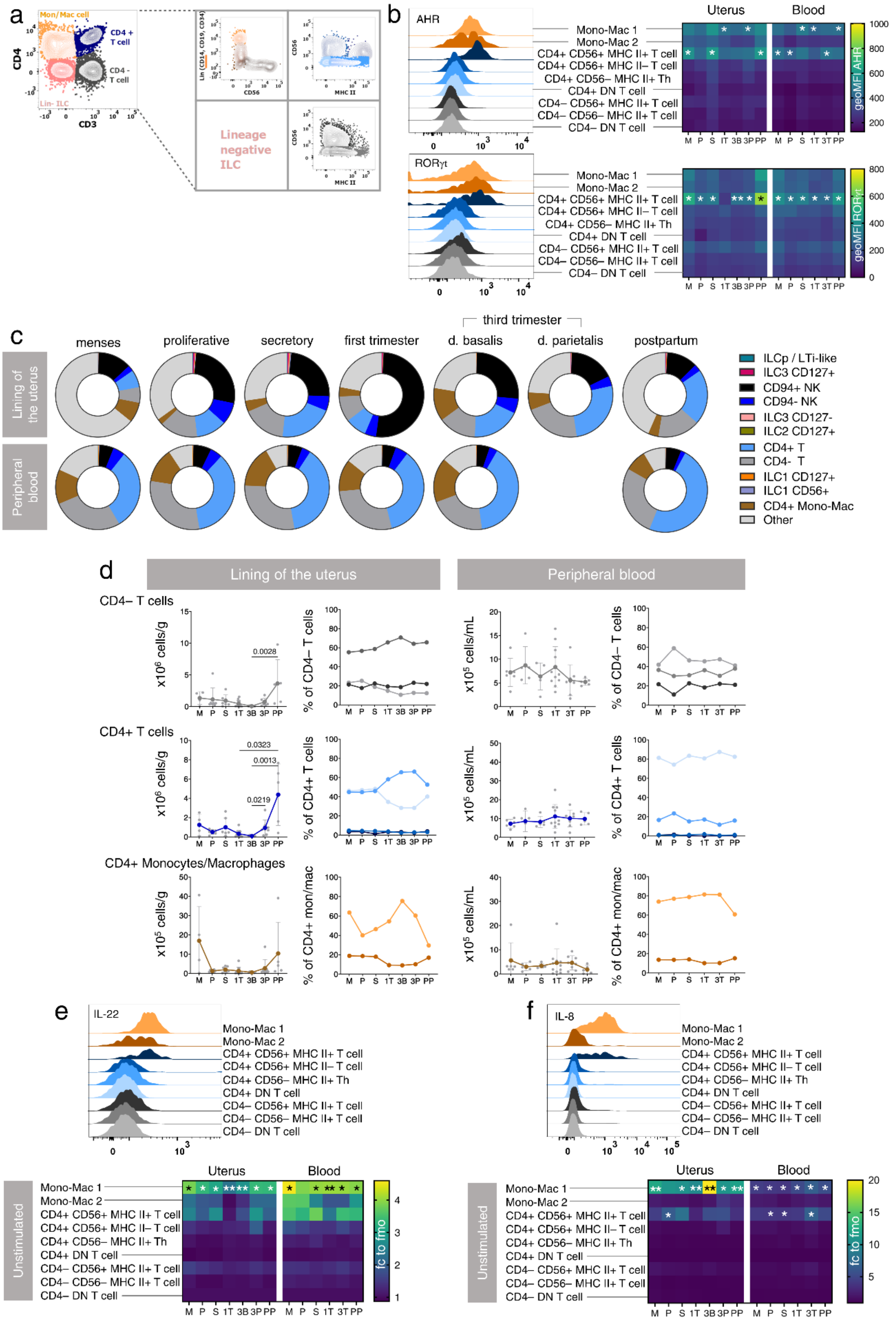


**Supplementary Figure 2.** Immune cells other than ILCs expressing IL-22 and IL-8 at the human uterine mucosal interface. **a** Flow cytometric gating strategy for non-ILC immune cells. **b** Expression of transcription factors AHR and RORγt by non-ILC immune cells in both the uterus and blood. **c** Representative proportion of a broad spectrum of immune cells in the uterus and blood in different reproductive stages. **d** Absolute number of non-ILC immune cells in the uterus and blood at any given stage of a healthy reproductive cycle. Proportions of simple non-ILC subpopulations are shown. **e** Intracellular IL-22 production by non-ILC immune cells. **f** Intracellular IL-8 production by non-ILC immune cells. Significance was determined via Z score transformation (GraphPad Prism).

a

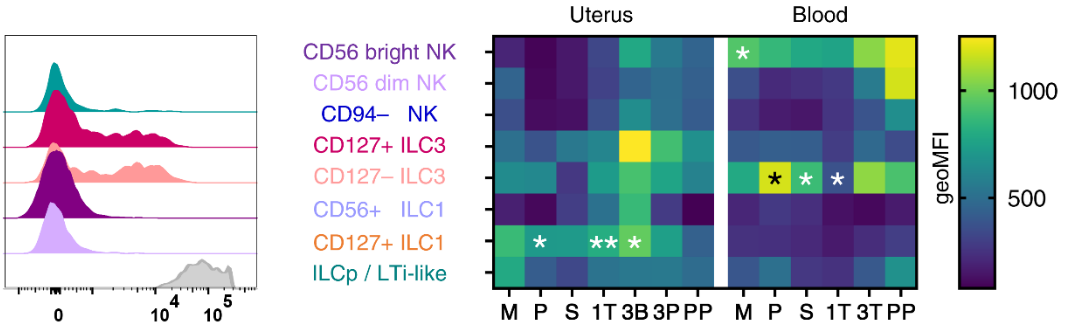


**Supplementary Figure 3. a** Histogram and heatmap representation of MHC II expression by uterine and peripheral blood ILC3. Z score transformed statistics (one-tailed), * $p < 0.05$, ** $p < 0.01$.

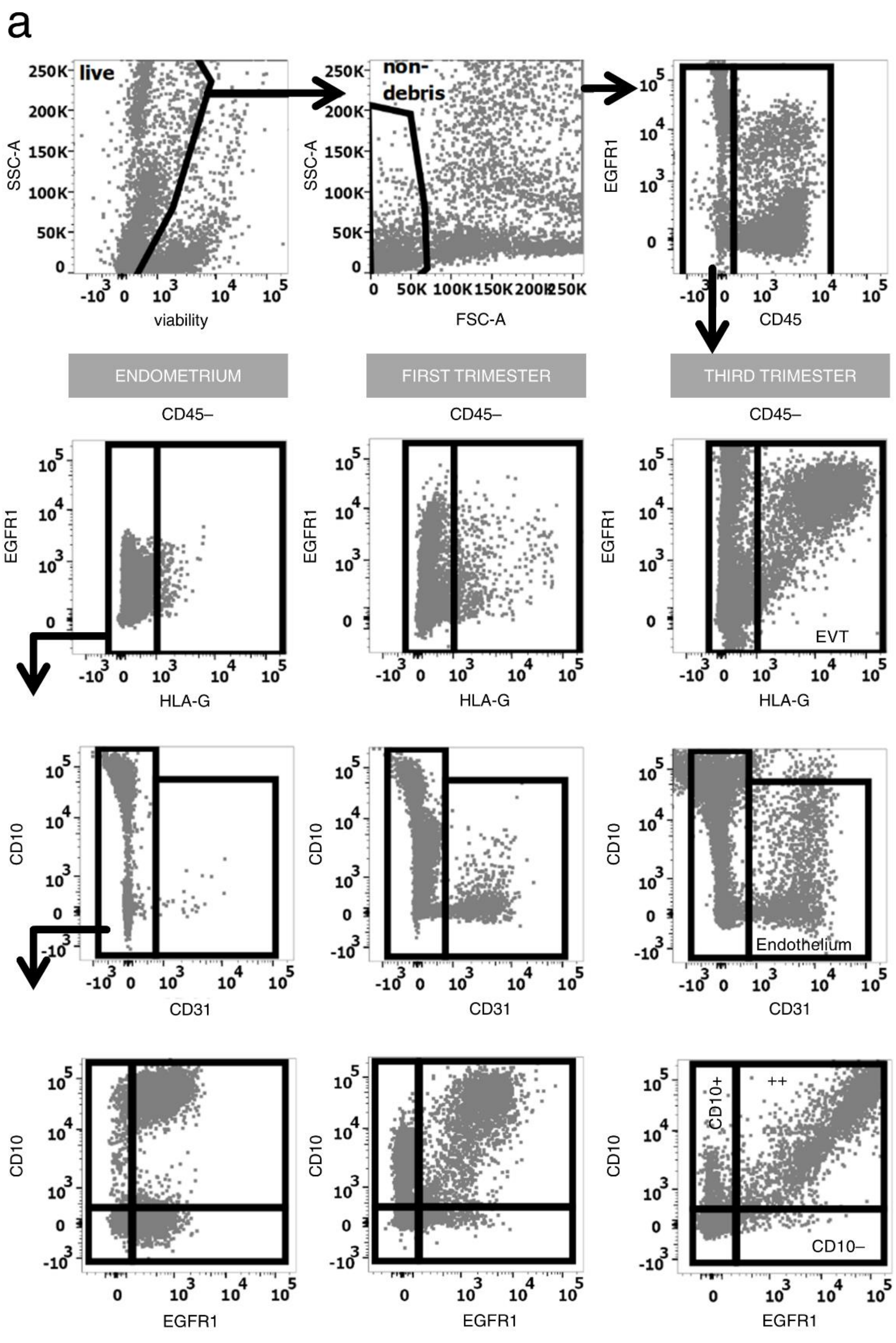


**Supplementary Figure 4. a** Complete flow cytometric gating strategy for non-immune cells from the lining of the uterus.

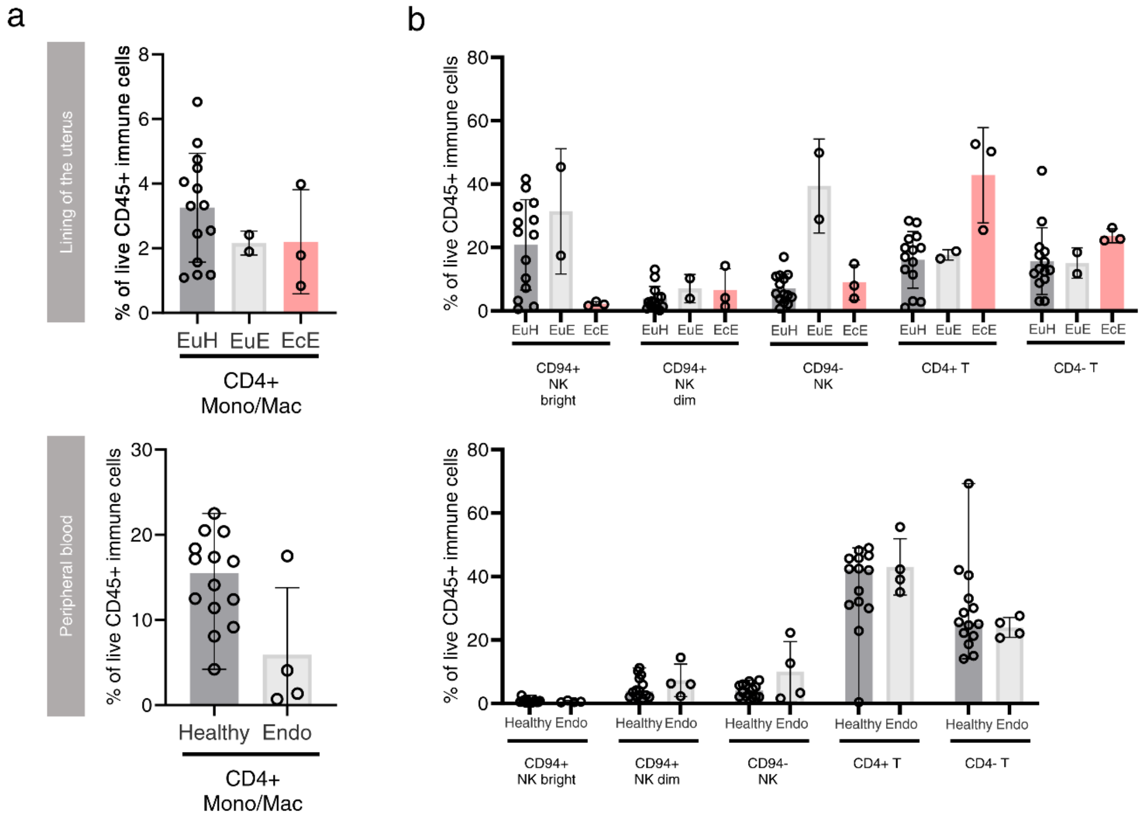


**Supplementary Figure 5. a** Frequency of monocytes/macrophages in the uterus and blood of donors with endometriosis relative to the normal baseline. **b** Frequency of NK cells and subsets of T cells in the uterus and blood of donors with endometriosis relative to the normal baseline.
