## Supplementary_Tables for "Dynamic roles of ILC3 in endometrial repair and regeneration"

Supplementary Table 1. Donor demographic information for eutopic endometrial samples taken from healthy non-pregnant women at different phases of the menstrual cycle.

| Sample ID | Age | Parity | Cycle day | Cycle length | Serum progesterone (nmol/L) | Phase assigned | Assessment | Comorbidity / Medication | Self-assigned ethnicity |
|---|---|---|---|---|---|---|---|---|---|
| E18 | 27 | nulliparous | 10 | 30 | 1 | Proliferative | Phenotype | None | Caucasian |
| E20 | 24 | nulliparous | 24 | 25 | 47 | Secretory | Phenotype | None | Caucasian |
| E22 | 29 | nulliparous | 7 | 28 | <1 | Proliferative | Phenotype | None | Caucasian |
| E24 | 32 | multiparous | n/a | n/a | <1 | Postpartum | Phenotype | None | Indian |
| E25 | 29 | nulliparous | 20 | 35 | 7 | Secretory | Phenotype | None | Caucasian |
| E26 | 36 | multiparous | 1 | 28 | 3 | Menses | Phenotype | None | Caucasian |
| E27 | 25 | nulliparous | 13 | 21 | 5 | Secretory | Phenotype | None | Caucasian |
| E28 | 23 | nulliparous | 2 | 28 | 1 | Menses | Phenotype | None | Caucasian |
| E36 | 23 | nulliparous | 1 | 29 | 1 | Menses | Phenotype | None | Afro-Caribbean |
| E37 | 40 | multiparous | 21 | 28 | 28 | Secretory | Phenotype | None | Caucasian |
| E38 | 24 | nulliparous | 17 | 34 | 6 | Secretory | Phenotype | None | Caucasian |
| E41 | 25 | nulliparous | 17 | 28 | NT | Secretory | Function | 4 months COCP withdrawn | Caucasian |
| E43 | 33 | multiparous | 19 | 28 | 22 | Secretory | Function | none | Caucasian |
| E44 | 38 | multiparous | 1 | 32 | 1 | Menses | Phenotype / Function | multiple miscarriages (x7) | Caucasian |
| E45 | 24 | nulliparous | 19 | 28 | 8 | Secretory | Phenotype | sertraline, lamotrigine, pregabalin | Caucasian |
| E47 | 31 | multiparous | 10 | 28 | 5 | Proliferative | Phenotype | None | Afro-Caribbean |

| | | | | | | | | | |
|---|---|---|---|---|---|---|---|---|---|
| E48 | 23 | nulliparous | 8 | 28 | <1 | Proliferative | Phenotype | 6 months COCP withdrawn | Caucasian |
| E49 | 32 | multiparous | 28 | 31 | 8 | Secretory | Function | Breastfeeding | Caucasian |
| E50 | 19 | nulliparous | 27 | 28 | 20 | Secretory | Function | None | Caucasian |
| E52 | 32 | nulliparous | 17 | 28 | 13 | Secretory | Function | None | Afro-Caribbean |
| E53 | 40 | nulliparous | 11 | 28 | 1 | Proliferative | Phenotype | Recent UTI: (completed Trimethoprim), covid-vaccinated 2 weeks prior | Caucasian |
| E56 | 36 | multiparous | 26 | 30 | 32 | Secretory | Phenotype | None | Pakistan |
| E57 | 37 | multiparous | n/a | n/a | 3 | Postpartum | Phenotype | None | North African |
| E59 | 29 | multiparous | n/a | n/a | FT | Postpartum | Phenotype | None | Pakistan |
| E61 | 22 | nulliparous | 17 | 28 | 1 | Secretory | Receptor | None, low serum progesterone | Caucasian |
| E62 | 33 | nulliparous | 27 | 30 | 34 | Secretory | Receptor | None | Caucasian |
| E63 | 23 | nulliparous | 7 | 36 | NT | Proliferative | Function / Receptor | None | Afro-Caribbean |
| E64 | 32 | nulliparous | 28 | 28 | 14 | Secretory | Receptor | Spironolactone for Acne | Caucasian |
| E65 | 28 | multiparous | n/a | n/a | <1 | Postpartum | Function / Receptor | Nonee | Asian - Other |
| E66 | 24 | nulliparous | 7 | 28 | 1 | Proliferative | Function | None | Caucasian |
| E68 | 36 | multiparous | n/a | n/a | NT | Postpartum | Phenotype / Function / Receptor | None | Afro-Caribbean |
| E70 | 34 | multiparous | 13 | 28 | 8 | Proliferative | Phenotype / Function / Receptor | None | Caucasian |
| E73 | 34 | nulliparous | 8 | 28 | <1 | Proliferative | Phenotype / Function | High BMI, previous herpes | Caucasian |
| E74 | 24 | nulliparous | 1 | 28 | 1 | Menses | Phenotype / Function | Anti-depressants | Asian - Other |
| E75 | 27 | nulliparous | 7 | 28 | <1.6 | Proliferative | Function | None | Asian |
| E76 | 26 | nulliparous | 4 | 28 | <1.6 | Menses | Phenotype / Function | None | Caucasian |

| | | | | | | | | | |
|---|---|---|---|---|---|---|---|---|---|
| E77 | 21 | nulliparous | 29 | 35 | 1.6 | Secretory | Receptor | Chronic pain, long cycle (query PCOS), low serum progesterone | Caucasian |
| E79 | 23 | nulliparous | 1 | 28 | NT | Menses | Function | None | Caucasian |
| E80 | 25 | nulliparous | 5 | 28 | NT | Menses | Phenotype / Function | None | Caucasian |
| E83 | 24 | nulliparous | 31 | 31 | NT | Secretory | Function | None | Caucasian |
| E84 | 30 | nulliparous | 10 | 28 | NT | Proliferative | Function | None | Other |
| E85 | 33 | multiparous | n/a | n/a | NT | Postpartum | Phenotype / Function | None | Caucasian |

Key

Parity: – indicates if patient has had a previous livebirth (birth of a live offspring > 24 weeks of gestation). Nulliparous refers to patients who were not pregnant at point of samples collection and had no previous livebirths, and this may include previous miscarriages or termination of pregnancies. Primiparous refers to patients who were pregnant at point of sample collection but had no previous livebirths, and this may include previous miscarriages and termination of pregnancies. Multiparous refers to patients who had previous livebirths.

NT – not tested; FT – failed test (irrelevant for assigned samples where values are expected to be low or negative).

PCOS – polycystic ovarian syndrome

BMI – body mass index

COCP – combined oral contraceptive pill

UTI – urinary tract infection

Supplementary 2. Donor demographic information for eutopic endometrial samples and ectopic lesions taken from women with endometriosis.

| sample ID | Age | Parity | Cycle day | Cycle length | Serum progesterone (nmol/L) | Phase assigned | Assessment | Tissue type | Comorbidity / Medication | Self-assigned ethnicity |
|---|---|---|---|---|---|---|---|---|---|---|
| E34 | 32 | nulliparous | 14 | 28 | 38 | Secretory | Phenotype / Function | Eutopic endometrium | none | Caucasian |
| E58 | 34 | nulliparous | 6 | 28 | <1 | Proliferative | Phenotype | Eutopic endometrium | none | Caucasian |
| ACE01 | 27 | nulliparous | 22 | 30 | NT | Secretory | Phenotype | Ectopic ovarian endometrioma (OE) | none | Middle eastern |
| ACE02 | 34 | nulliparous | n/a | n/a | NT | Contraceptive altered | Phenotype | Ectopic peritoneal nodule (NO) | Patient on continuous combined oral contraceptive pill | Middle eastern |
| ACE04 | 31 | nulliparous | 7 | 30 | NT | Proliferative | Phenotype / Function | Ectopic ovarian endometrioma (OE) | Infertility (having had five failed previous embryo transfers) | Caucasian |
| ACE05 | 45 | nulliparous | 32 | 32 | NT | Late Secretory | Phenotype | Ectopic ovarian endometrioma (OE) | none | Caucasian |

Supplementary Table 3. Donor demographic information for eutopic endometrial samples from the EXPPECT biobank (University of Edinburgh). Samples were collected with informed consent from patients with endometriosis of known fertility status, and controls without endometriosis. Controls may have a benign gynecological condition, such as pain, also with confirmed fertility status. Samples are derived from women of reproductive age, fixed in 4% paraformaldehyde, dehydrated in 70% ethanol and embedded into paraffin wax blocks for storage.

| sample ID | Tissue type | Endometriosis stage: revised American Society for Reproductive Medicine (rASRM) classification | Fertility status checked | Phase assigned | Assessment | Biobank year |
|---|---|---|---|---|---|---|
| 1083 | Eutopic endometrium_Control | na | fertile | Early Secretory | Immunofluorescence | 2016 |
| 3266 | Eutopic endometrium_Control | na | fertile | Early Secretory | Immunofluorescence | 2016 |
| 4022 | Eutopic endometrium_Control | na | fertile | Mid-Secretory | Immunofluorescence | 2017 |
| 1076 | Eutopic endometrium_Endometriosis | Stage 4 (>40)- Severe | fertile | Early Secretory | Immunofluorescence | 2016 |
| 1033 | Eutopic endometrium_Endometriosis | Stage 4 (>40)- Severe | fertile | Early Secretory | Immunofluorescence | 2017 |
| 1805 | Eutopic endometrium_Endometriosis | Stage 4 (>40)- Severe | fertile | Mid-Secretory | Immunofluorescence | 2019 |
| 3267 | Eutopic endometrium_Endometriosis | Stage 4 (>40)- Severe | infertile | Early Secretory | Immunofluorescence | 2016 |
| 1658 | Eutopic endometrium_Endometriosis | Stage 4 (>40)- Severe | infertile | Early Secretory | Immunofluorescence | 2017 |
| 1262 | Eutopic endometrium_Endometriosis | Stage 4 (>40)- Severe | infertile | Late Secretory | Immunofluorescence | 2016 |

Supplementary Table 4. Donor demographic information for decidual samples taken from women during different pregnancy stages.

| sample ID | Age | Parity | Gestation | Reproductive Stage | Delivery | Assessment | Pregnancy-related Comorbidity / Medication | Self-assigned ethnicity |
|---|---|---|---|---|---|---|---|---|
| TOP10 | 24 | Multiparous | 8+6 | 1T | Personally-requested termination | Phenotype / Function | None | Caucasian |
| TOP12 | 20 | Primiparous | 6+6 | 1T | Personally-requested termination | Phenotype / Function / Receptor | None | Caucasian |
| TOP13 | 32 | Multiparous | 10+4 | 1T | Personally-requested termination | Phenotype / Function / Receptor | high BMI | Caucasian |
| TOP14 | 25 | Primiparous | 11+5 | 1T | Personally-requested termination | Phenotype / Function / Receptor | None | Asian_Indian |
| TOP15 | 22 | Primiparous | 8+3 | 1T | Personally-requested termination | Phenotype / Function / Receptor | Cannabis use, chronic ITP | Caucasian |
| TOP16 | 21 | Multiparous | 13/0 | 1T | Personally-requested termination | Phenotype / Function / Receptor | high BMI | Caucasian |
| TOP17 | 33 | Multiparous | 12/6 | 1T | Personally-requested termination | Phenotype / Function / Receptor | None | Caucasian |
| TOP19 | 30 | Multiparous | 6/0 | 1T | Personally-requested termination | Phenotype / Function | High BMI | Caucasian |
| TOP21 | 40 | Multiparous | 8/6 | 1T | Personally-requested termination | Phenotype / Function | On allopurinol (self-medicating) | Asian_Indian |
| TOP22 | 40 | Multiparous | 10+5 | 1T | Personally-requested termination | Phenotype / Function | SLE | Mixed |
| TOP23 | 26 | Multiparous | 6/0 | 1T | Personally-requested termination | Phenotype / Function | Cervical dyskaryosis (CIN 2) | Caucasian |
| TOP24 | 34 | Multiparous | 9+6 | 1T | Personally-requested termination | Phenotype / Function | asthma | Caucasian |
| TOP25 | 19 | Primiparous | 10+0 | 1T | Personally-requested termination | Phenotype / Function | Depression on sertraline and circadin | Caucasian |

| | | | | | | | | |
|---|---|---|---|---|---|---|---|---|
| AC_TNL01 | 35 | Multiparous | 37+0 | 3T | Elective caesarean | Phenotype/Receptors | Pregnancy induced hypertension | White – Any other White background |
| AC_TNL02 | 30 | Primiparous | 38+0 | 3T | Elective caesarean | Phenotype | Ulcerative colitis (Mesalazine), bile acid malabsorption (cholestagel) | White – British |
| AC_TNL03 | 39 | Multiparous | 39+0 | 3T | Elective caesarean | Phenotype | none | White – Any other White background |
| AC_TNL04 | 35 | Multiparous | 39+0 | 3T | Elective caesarean | Function | none | Any Other Ethnic Group |
| AC_TNL05 | 35 | Primiparous | 39+0 | 3T | Elective caesarean | Phenotype | PCOS | White – British |
| AC_TNL06 | 40 | Primiparous | 38+6 | 3T | Elective caesarean | Phenotype | none | |
| AC_TNL07 | 41 | Multiparous | 39+3 | 3T | Elective caesarean | Phenotype | query hypothyroidism; loop exciscion of cervix to remove pre-cancerous cells | White other (Eastern European) |
| AC_TNL09 | 34 | Primiparous | 39+0 | 3T | Elective caesarean | Function / Receptors | hypothyroidism | Caucasian |
| AC_TNL10 | 41 | Primiparous | 39+1 | 3T | Elective caesarean | Receptors | Ha’shimoto's thyroiditis; loop excision of cervix to remove pre-cancerous cells | White British |
| AC_TNL11 | 36 | Primiparous | 39+0 | | Elective caesarean | Function / Receptors | previous shoulder operation | White British |
| AC_TNL13 | 31 | Multiparous | 39+0 | 3T | Elective caesarean | Function | Aortic coarctation | White British |
| AC_TNL14 | 41 | Primiparous | 39+0 | 3T | Elective caesarean | Phenotype / Function | Aspirin, cyclogest | White – Any other White background |
| AC_TNL15 | 41 | Primiparous | 39+3 | 3T | Elective caesarean | Phenotype / Function | none | Any Other Ethnic Group |

Key – as above.

Supplementary Table 5. Donor sample demographic information for decidual samples taken from women having pregnancy complications during different pregnancy stages.

| sample ID | Age | Parity | Gestation | Reproductive Stage | Delivery | Assessment | Pregnancy-related Comorbidity / Medication | Self-assigned ethnicity |
|---|---|---|---|---|---|---|---|---|
| PTNL01 (PPROM) | 36 | Multiparous | 27/2 | 2T | Emergency caesarean | Phenotype / Function | preterm chorioamnionitis (chorio) | E~~u~~ropean - Albanian |
| PTNL02 (preeclampsia) | 45 | Multiparous | 24/4 | 2T | Emergency caesarean | Phenotype / Function | preterm preeclamptic | Black or Black British-African |
| PTNL03 (preeclampsia) | 37 | Primiparous | 25/5 | 2T | Emergency caesarean | Phenotype / Function | preterm preeclamptic | White – British |
| PTNL04 (preeclampsia) | 28 | Primiparous | 27/0 | 2T | Emergency caesarean | Phenotype / Function | preterm preeclamptic | Other – not stated |
| PTNL05 (post-congenital CMV) | 32 | Multiparous | 26/6 | 2T | Emergency caesarean | Phenotype / Function | preterm NO-PET/Bact. Infection (but CMV) | Mid-~~E~~astern - Afghan |
| PTNL06 (preeclampsia) | 28 | Primiparous | 30/6 | 3T | Scheduled caesarean | Phenotype / Function | third trimester preeclampti~~c~~ | White - British |
| CTNL01 (preeclampsia) | 48 | Primiparous | 37 | 3T | Scheduled caesarean | Phenotype / Function | third trimester preeclampitic | White – Any other white background |
| CTNL02 (preeclampsia) | 33 | Primiparous | 36/5 | 3T | Scheduled caesarean | Phenotype / Function | third trimester preeclampitic | White – Any other white background |
| CTL01 (PPROM) | 43 | Multiparous | 37 | 3T | Induced vaginal | Phenotype / Function | term no chorio (PPROM 20/6~~)~~ | White - British |
| CTL02 (septic in labour) | 32 | Primiparous | 41/1 | 3T | Emergency caesarean | Phenotype / Function | septic in labour (blunt infection) | South-European |

Supplementary Table 6. List of fluorescent-conjugated antibodies used in flow cytometry experiments.

| **Antibody** | **Clone** | **Fluorophore** | **Dilution** | **Host animal** | **Manufacturer** | **Catalog #** | **RRID** |
|---|---|---|---|---|---|---|---|
| **ILC3 backbone panel** | | | | | | | |
| CD3 | SK7 | APC-eFluor 780 | 1/200 | Ms | eBioscience | 47-0036 | AB_10718679 |
| CD4 | OKT4 | Brilliant Violet 785 | 1/100 | Ms | Biolegend | 317441 | AB_2561365 |
| ~~CD7~~ | ~~CD7-6B7~~ | ~~PerCP-Cy5.5~~ | ~~1/100~~ | ~~Ms~~ | ~~Biolegend~~ | ~~343115~~ | ~~AB_2632911~~ |
| CD14 | 63D3 | FITC | 1/100 | Ms | Biolegend | 367115 | AB_2571928 |
| CD19 | HIB19 | FITC | 1/100 | Ms | Biolegend | 302205 | AB_314235 |
| CD34 | 561 | FITC | 1/100 | Ms | Biolegend | 343603 | AB_1732030 |
| CD45 | 2D1 | Alexa Fluor 700 | 1/100 | Ms | Biolegend | 368513 | AB_2566373 |
| CD56 | NCAM16.2 | Brilliant Violet 510 | 1/200 | Ms | BD Biosciences | 563041 | AB_2732786 |
| CD94 | DX22 | PerCP-Cy5.5 | 1/200 | Ms | Biolegend | 305514 | AB_2565522 |
| CD117 | 104D2 | PE-Dazzle 594 | 1/200 | Ms | Biolegend | 313225 | AB_2566212 |
| CD127 | HIL-7R-M21 | Alexa Fluor 647 | 1/20 | Ms | BD Biosciences | 558598 | AB_647113 |
| CD294 (CRTH2) | BM16 | Brilliant Violet 605 | 1/50 | Rat | Biolegend | 350121 | AB_2566759 |
| HLA-DR,DQ,DP | Tu39 | Brilliant Violet 711 | 1/100 | Ms | BD Biosciences | 740784 | AB_2740447 |
| **ILC phenotypic transcription factors** | | | | | | | |
| RORγt | Q21-559 | Brilliant Violet 650 | 1/20 | Ms | BD Biosciences | 563424 | AB_2738197 |
| AHR | FF3399 | PE | 1/20 | Ms | eBioscience | 12-9854 | AB_2572745 |
| GATA3 | L50-823 | PE-Cy7 | 1/25 | Ms | BD Biosciences | 560405 | AB_1645544 |
| **ILC3 functional effector molecules** | | | | | | | |
| IL-8 | G265-8 | PE | 1/100 | Ms | BD Biosciences | 554720 | AB_395529 |
| IL-22 | 22URTI | PE-Cy7 | 1/20 | Ms | eBioscience | 25-7229 | AB_10853659 |
| **Effector molecule receptor panel** | | | | | | | |
| CD10 | HI10a | Brilliant Violet 785 | 1/100 | Ms | Biolegend | 312237 | AB_2860829 |
| CD31 (PECAM-1) | WM-59 (WM59) | APC-eFluor 780 | 1/100 | Ms | eBioscience | 47-0319 | AB_10730582 |
| CXCR1 | 5A12 | Brilliant Violet 711 | 1/100 | Ms | BD Biosciences | 743423 | AB_2741496 |
| CXCR2 | 5E8/CXCR2 | PE-Dazzle 594 | 1/100 | Ms | Biolegend | 320722 | AB_2750215 |
| EGFR1 | AY13 | PE-Cy7 | 1/200 | Ms | Biolegend | 352910 | AB_2562159 |
| HLA-G | MEM/9 | FITC | 1/25 | Ms | Invitrogen | MA1-19591 | AB_1076722 |
| IL10Rβ | 90220 | Alexa Fluor 647 | 1/20 | Ms | R&D Systems | FAB874R-100UG | na |
| IL22Rα1 | 305405 | PE | 1/20 | Ms | R&D Systems | FAB2770P | AB_2124369 |

**Supplementary Table 7.a. Origin and amounts of primary antibodies used in fluorescent immunohistochemistry.**

| Antibody | Clone | Conjugation | Dilution | Origin | Manufacturer | Retrieval solution | Catalog # | RRID |
|---|---|---|---|---|---|---|---|---|
| Anti-human CD117 (c-Kit) | YR145 | unconjugated | 1/200 | rabbit | Abcam | Tris-EDTA pH9.0 | ab32363 | AB_731513 |
| Anti-human CD56 | 123C3 | unconjugated | 1/500 | mouse | Invitrogen | EDTA pH8.0 | 18–0152 | AB_138599 |
| Anti-human CD31 | C31.3 + JC/70A | unconjugated | 1/200 | mouse | Abcam | Citrate pH6.0 | ab199012 | AB_2756834; AB_307284 |
| Anti-human IL-22 | polyclonal | unconjugated | 1/50 | goat | R&D Systems | Citrate pH6.0 | AF782 | AB_355597 |

**Supplementary Table 7.b. Origin and amounts of secondary antibodies used in fluorescent immunohistochemistry.**

| Antibody | Clone | Conjugation | Dilution | Origin | Manufacturer | Catalog # | RRID |
|---|---|---|---|---|---|---|---|
| Anti-goat IgG | polyclonal | HRP | 1/500 | donkey | Santa Cruz | sc-2020 | AB_631728 |
| Anti-rabbit IgG | polyclonal | HRP | 1/500 | donkey | Jackson Immuno Research | 711-036-152 | AB_2340590 |
| Anti-mouse IgG | polyclonal | HRP | 1/500 | donkey | Jackson Immuno Research | 715-036-150 | AB_2340773 |

**Supplementary Table 7.c. Fluorescent dyes used for marker detection under catalysis of horse-radish peroxidase.**

| Antibody | Manufacturer | Catalog # |
|---|---|---|
| Opal 520 (green) | Akoya Biosciences | FP1487001KT |
| Opal 570 (red) | Akoya Biosciences | FP1488001KT |
| Opal 650 (blue) | Akoya Biosciences | FP1496001KT |
| 1X Diluent | Akoya Biosciences | FP1498 |
